# A landscape of differentiated biological processes involved in the initiation of sex differentiation in zebrafish

**DOI:** 10.1101/2022.04.14.488425

**Authors:** Ding Ye, Yi-Xuan Tu, Houpeng Wang, Mudan He, Yaqing Wang, Zhengfang Chen, Zhen-Xia Chen, Yonghua Sun

## Abstract

Zebrafish (*Danio rerio*) has been used as a promising animal model to study gonadal development and gametogenesis. Although previous studies have identified critical molecules participating in zebrafish gonad differentiation, a landscape view of the biological processes involved in this process is still lacking. Here we isolated intact zebrafish differentiating gonads, at 25 days post-fertilization (dpf) and 30 dpf, and conducted RNA-seq analysis between the juvenile gonads that tended to develop into ovaries or testes. Our study demonstrates that the juvenile ovary and testis at 25 dpf and 30 dpf are different at the level of biological process. During ovary differentiation, the biological processes related to metabolic activities in production of energy and maternal substances, RNA degradation, and DNA repair were enriched. During testis differentiation, the biological processes related to cell proliferation, differentiation, and morphogenesis were enriched, with a total of 15 signaling pathways. Notably, we reveal that the immune-related processes are extensively involved in the regulation of testis development. Overall, this study provides a landscape of differentiated biological processes and novel insights into the initiation of sex differentiation in zebrafish.

## 1. Introduction

Gonad differentiation is a complicated process that a bio-potential gonad decides to develop to an ovary or a testis. Zebrafish (*Danio rerio*) has been used as an animal model to study gonadal differentiation and gametogenesis (Zhu and Ge 2018, Li and Ge 2020), and a promising model for genome editing and directional breeding of farmed fishes (Ye, Zhu et al. 2015, Sun 2017, Sun and Zhu 2019). Recently, genome-edited sperms from a different subfamily were produced in zebrafish testes by spermatogonial stem cell transplantation, which opens an era for surrogate reproduction of farmed fishes in the future (Zhang, Hao et al. 2021). In zebrafish, it is generally accepted that the bi-potential gonads start to differentiate toward to an ovary or a testis during 20 dpf to 25 dpf (Chen, Liu et al. 2017, Kossack and Draper 2019). From about 25 dpf to 35 pdf, ovary and testis increasingly differ in morphology as gonad differentiation proceeding. In the presumptive ovary, the oocytes continue to grow and the gonadal size increases dramatically (Uchida, Yamashita et al. 2002, Ye, Zhu et al. 2019). In contrast, in the presumptive testis, the gonadal size mildly increases and a certain level of cell apoptosis occurs (Uchida, Yamashita et al. 2002, Ye, Zhu et al. 2019). Recently, we realized the life-time labeling of zebrafish germline with a transgenic line, *Tg*(*piwil1-egfp-nos3UTR*)^*ihb327Tg*^ (Ye, Zhu et al. 2019, Ye and Sun 2021), and revealed that the presumptive female zebrafish generally have more primordial germ cells (PGCs) at embryonic stage and relatively large gonads at juvenile stage, while the presumptive males have less PGCs and juvenile gonads with small size (Ye, Zhu et al. 2019).

Zebrafish gonad differentiation is regulated by multiple genes, accompanied with dramatic difference between the presumptive ovary and the presumptive testis at cellular and molecular level. There are several cellular events happening during gonad differentiation, such as mitosis of germline stem cells and gonad somatic cells, meiosis of differentiating germ cells, and cell apoptosis in juvenile testis (Kossack and Draper 2019). Steroid biosynthetic genes (*cyp19a1a* (Lau, Zhang et al. 2016), *cyp11c1* (Zhang, Ye et al. 2020), *cyp11a1* (Tong, Hsu et al. 2010), *cyp11a2* (Wang, Ye et al. 2021), *star* (Shang, Peng et al. 2019)), receptor-coding genes (*androgen receptor* (Crowder, Lassiter et al. 2018, Tang, Chen et al. 2018, Yu, Zhang et al. 2018), *estrogen receptor* (Lu, Cui et al. 2017, Chen, Tang et al. 2018), *fshr* (Zhang, Lau et al. 2015), *bmpr1bb* (Little and Mullins 2009)), transcription factors (*dmrt1* (Lin, Mei et al. 2017, Webster, Schach et al. 2017, Romano, Kaufman et al. 2020)), signaling molecules (*gdf9* (Chen, Liu et al. 2017), *inhbab* (Wang and Ge 2004, DiMuccio, Mukai et al. 2005), *inhbb* (Wang and Ge 2004), *bmp15* (Dranow, Hu et al. 2016), *wnt4* (Kossack, High et al. 2019), *fgf24* (Leerberg, Sano et al. 2017), *amh* (Lin, Mei et al. 2017, Zhang, Zhu et al. 2020)) have been reported to be critical for gonad differentiation and gametogenesis in zebrafish.

In recent years, several genome-wide transcriptional analyses of zebrafish gonads were conducted with mature gonads or gonad-containing body pieces. By isolating mature gonads at adult stage and trunk regions containing juvenile gonads at 14 dpf and 22 dpf, the sexually dimorphic expression profiles in both sexes were analyzed by micro-array (Sreenivasan, Cai et al. 2008, Tzung, Goto et al. 2015). By using whole juvenile fish at 25 dpf, the transcriptomic change during sex differentiation was also studied through RNA sequencing (Chen, Wang et al. 2020). However, to better understand the gene expression profile and biological events at the onset of gonad differentiation, it is required to use isolated intact juvenile gonads that start to differentiate as the samples for RNA sequencing.

Here in this study, by using the transgenic zebrafish *Tg*(*piwil1-egfp-nos3UTR*)^*ihb327Tg*^, we were able to isolate intact zebrafish juvenile ovaries and testes at 25 dpf and 30 dpf, and conducted RNA-seq analysis. Our study demonstrates that the juvenile ovaries and testes display significant differences at the level of biological process, and provides a landscape of differentiated biological processes and novel insights into the initiation of sex differentiation. Especially, the immune-related processes are suggested to be extensively involved in the regulation of testis development.

## 2. Materials and Methods

### 2.1 Fish stocks and collection of gonad samples

The transgenic zebrafish *Tg(piwil1-egfp-nos3UTR)*^*ihb327Tg*^ from the China Zebrafish Resource Center, National Aquatic Biological Resource Center (CZRC/NABRC, Wuhan, China) was described previously (Ye, Zhu et al. 2019). By using this transgenic fish, the germline can be lifetime labeled with enhanced green fluorescent protein (EGFP), which would show the gonad under a fluorescent stereomicroscope and help with precise dissection. Fish were bred and maintained according to the Zebrafish Book (Westerfield 2000).

### 2.2 Collection of gonad samples

By using the transgenic zebrafish *Tg(piwil1-egfp-nos3UTR)*^*ihb327Tg*^, the embryos with high number of PGCs (PGC number >36 per embryo at 1 dpf) and those with low number of PGCs (PGC number <25 per embryo at 1 dpf) were selected, and raised up as two groups as previously described (Ye, Zhu et al. 2019)(Figure 1A). To avoid the developmental differences among different juveniles (Parichy, Elizondo et al. 2009), the fish with similar standard length were selected, and the average standard length of the selected 25 dpf and 30 dpf fishes were about 1.0 centimeters (cm) and 1.4 cm, respectively (Figure 1B). The transgenic zebrafish larvae were anesthetized in 0.2 mg/mL MS-222 and dissected under a fluorescent stereomicroscope to isolate intact gonads using eye scissors and tweezers with fine tips. At least 20 gonads from each group were dissected and imaged, and their sizes were measured using Fiji software (Schindelin, Arganda-Carreras et al. 2012). Each gonad was immediately immersed into RNA Stabilizer (Vazyme) to protect RNA from degradation. The 3 smallest gonads were chosen as the testis samples, and the 3 biggest ones were chosen as the ovary samples.

**Fig. 1.**
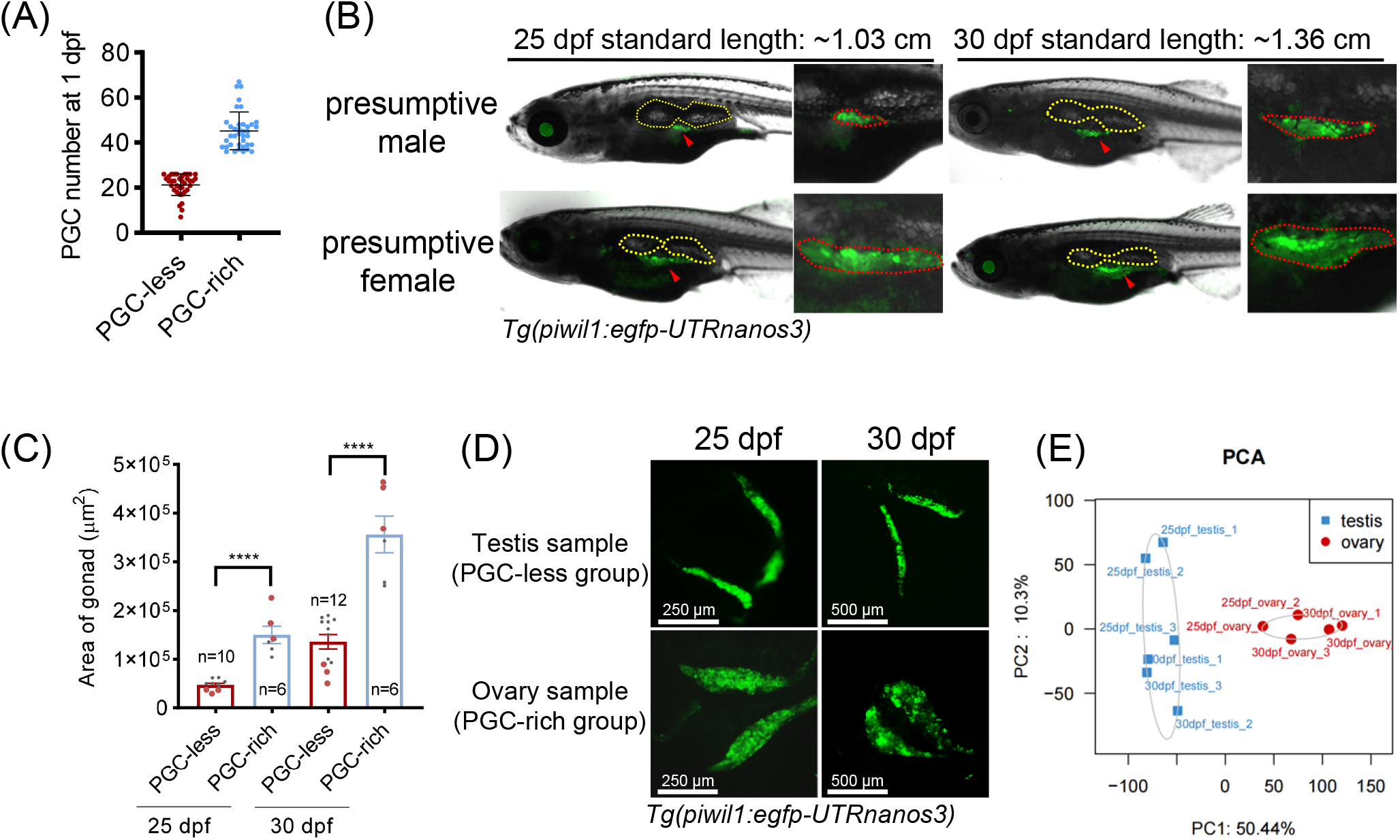
Study design, sample diagnose, PCA analysis and DEGs analysis. (A) The PGC number of the germline-labeled transgenic zebrafish embryos from selected PGC-less group and PGC-rich group at 1 dpf. (B) The representative images of the intact fish from the above two groups at 25 dpf and 30 dpf. The swim bladders are circled by dotted white lines, and the gonads are circled by dotted red lines. (C) The areas of the isolated gonads from the fish of the above two groups at 25 dpf and 30 dpf. The red dots represent the samples for RNA sequencing (n represent the fish number, “****” p<0.0001). (D) The representative images of the testis sample and the ovary sample from the 25 dpf and 30 dpf germline-labeled transgenic zebrafish which came from the PGC-less group and the PGC-rich group. (E) The PCA map of the gonad samples.

### 2.3 Preparation of RNA library and RNA sequencing

The cDNA Libraries with low-input RNAs were prepared according to manufacturer’s instructions. After washing off the RNA Stabilizer, RNA from an intact gonad was extracted and purified using the RNAprep pure Micro Kit (TIANGEN). The quality of RNA was determined by Nanodrop, Qubit and Labchip GX. For each sample, 5 ng of RNA was used for synthesis and amplification of cDNA (Discover-sc WTA Kit, N711, Vazyme), and the length distribution of cDNA was tested by Agilent2100. The cDNA libraries were then constructed using TruePrep DNA Library Prep Kit (D501, Vazyme). Sequencing was performed on Illumina Nextseq 500 with read length of 150 bp paired-end (PE) at the Analysis and Testing Center of Institute of Hydrobiology, Chinese Academy of Sciences, China. For each sample, about 6G clean data with Q30>90% were generated. The clean data were mapped to a zebrafish reference transcripts generated from genome GRCz11 Ensembl release 97 using Bowtie2 with default parameters (Langmead and Salzberg 2012). Transcripts were normalized and quantified by RSEM (RNA-Seq by Expectation Maximization) (Li and Dewey 2011). The principal component analysis (PCA) was performed using R package. Differential expression analysis was performed using DEseq2 (Love, Huber et al. 2014). Genes with the criteria |log2foldchange|>=1, adjusted P<0.05 were considered as differentially expressed genes (DEGs), and the DEGs were used for Gene Ontology (GO) enrichment analysis. The GO enrichment analysis and the related visualization were performed by ClusterProfiler 4.0 (Wu, Hu et al. 2021).

### 2.4 Validation of DEGs using quantitative PCR (qPCR)

Juvenile zebrafish at 30 dpf were collected to isolate gonads, and the sex types were identified as previously described [9]. Three ovaries or testes were combined, and the RNA was extracted. The RNA of 150 ng was reverse-transcribed using HiScript® II Reverse Transcriptase with random hexamers (Vazyme). Quantitative PCR (qPCR) was performed and the data was analyzed according to the MIQE (Minimum Information for Publication of Quantitative Real-Time PCR Experiments) guidelines (Bustin, Benes et al. 2009). *β-actin* was selected as the reference gene. All the qPCR gene-specific primers are listed in the Table S1.

## 3. Results and disscussion

### 3.1 Study design, data generation and quality control

In our previous study, we found that the variation on PGC number between the PGC-less and the PGC-rich groups persisted and led to different sex development, in which the PGC-less embryos tended to develop into males, while the PGC-rich embryos tended to develop into females (Ye, Zhu et al. 2019). Therefore, we utilized the *Tg(piwil1-egfp-nos3UTR)*^*ihb327Tg*^ embryos for the present study, and selected the PGC-less embryos (PGC number <25 per embryo at 1 dpf) and PGC-rich embryos (PGC number >36 per embryo at 1 dpf), and raised them in two groups. In accordance, at 25 dpf and 30 dpf, the juvenile gonads in the PGC-less group were significantly smaller than those from the PGC-rich group (Figure 1B, C). In order to screen the juvenile testes and juvenile ovaries at 25 dpf and 30 dpf for RNA-seq analysis, we isolated and selected the smallest intact gonads from the PGC-less group, and the biggest intact ones from the PGC-rich group for subsequent study (Figure 1D, See **Materials and Methods** for detail).

We performed 150 bp pair-ended sequencing of poly(A)+ RNA on single gonad with biological replicates from six 25 dpf and six 30 dpf intact gonads. For each gonad, an average of 46 million clean reads was generated. The average mean of Q20 and Q30 were 96% and 91%, respectively. A total of ∼46,000 transcripts and ∼25600 genes were detected in the gonad samples. To verify the gonad type once again, we performed PCA map using the FPKM expression matrix of all the samples, and the data showed that the testes and the ovaries were separated as two major clusters (Figure 1 E). These data indicate that the sexually dimorphic expression profiles start to be evident in the juvenile fish at as early as 25 dpf, which extends our understanding of sex differentiation of zebrafish.

### 3.2 Pro-female and pro-male genes

There were 2471 genes up-regulated in differentiating ovaries, whereas 2885 genes were down-regulated in differentiating testes (Figure 2A, Table S2). In the DEG list, some genes known to be involved in gonad development and/or gametogenesis were shown in the Figure 2 (Figure 2B, C) (Kossack and Draper 2019, Li and Ge 2020). Interestingly, *hsd3b1* and *hsd3b2*, both of which encode 3β-hydroxysteroid dehydrogenase, were differentially expressed in the testis or the ovary, suggesting that each paralog is under sex-specific regulation (Figure 2B, C; *hsd3b1* and *hsd3b2* are marked by star(*)). Genes coding both androgen receptor and estrogen receptor were highly expressed in the testis, implying both androgen and estrogen may play roles in testis development (Figure 2B). We also found sex biased transcription factors and signaling ligands which were differentially expressed in the testis or the ovary (Figure 2B, C). Moreover, genes encoding RNA binding proteins, *tdrd12* and *zar1*, were highly expressed in the ovary, while the thyroid activation gene *dio2* was highly expressed in the testis (Figure 2B, C). We selected some genes and validated their expressions by qPCR (Figure 2D). One of the selected gene *cyp11c1* has been found to promote testis differentiation in our recent study (Zhang, Ye et al. 2020). Another steroidgenic gene *cyp11a1* has been known to have dimorphic expression during early gonad differentiation, but the function has not been reported (Tong, Hsu et al. 2010). In our previous studies, we have generated the genetic mutant of *cyp11a2*, the zebrafish paralog of *cyp11a1*, and found the homozygotic mutant developing into all-male with hyperproliferative spermatogonia, and we proposed that *cyp11a1* may compensate for a partial function in the absence of *cyp11a2* in zebrafish (Wang, Ye et al. 2021). To study the exact role of *cyp11a1* in gonad development and gametogenesis, the double mutant of *cyp11a1* and *cyp11a2* would be much helpful. The enrichment of known pro-female and pro-male genes in the DEG list further supported that the sequenced samples were juvenile ovaries and juvenile testes with dimorphic expression patterns.

**Fig. 2.**
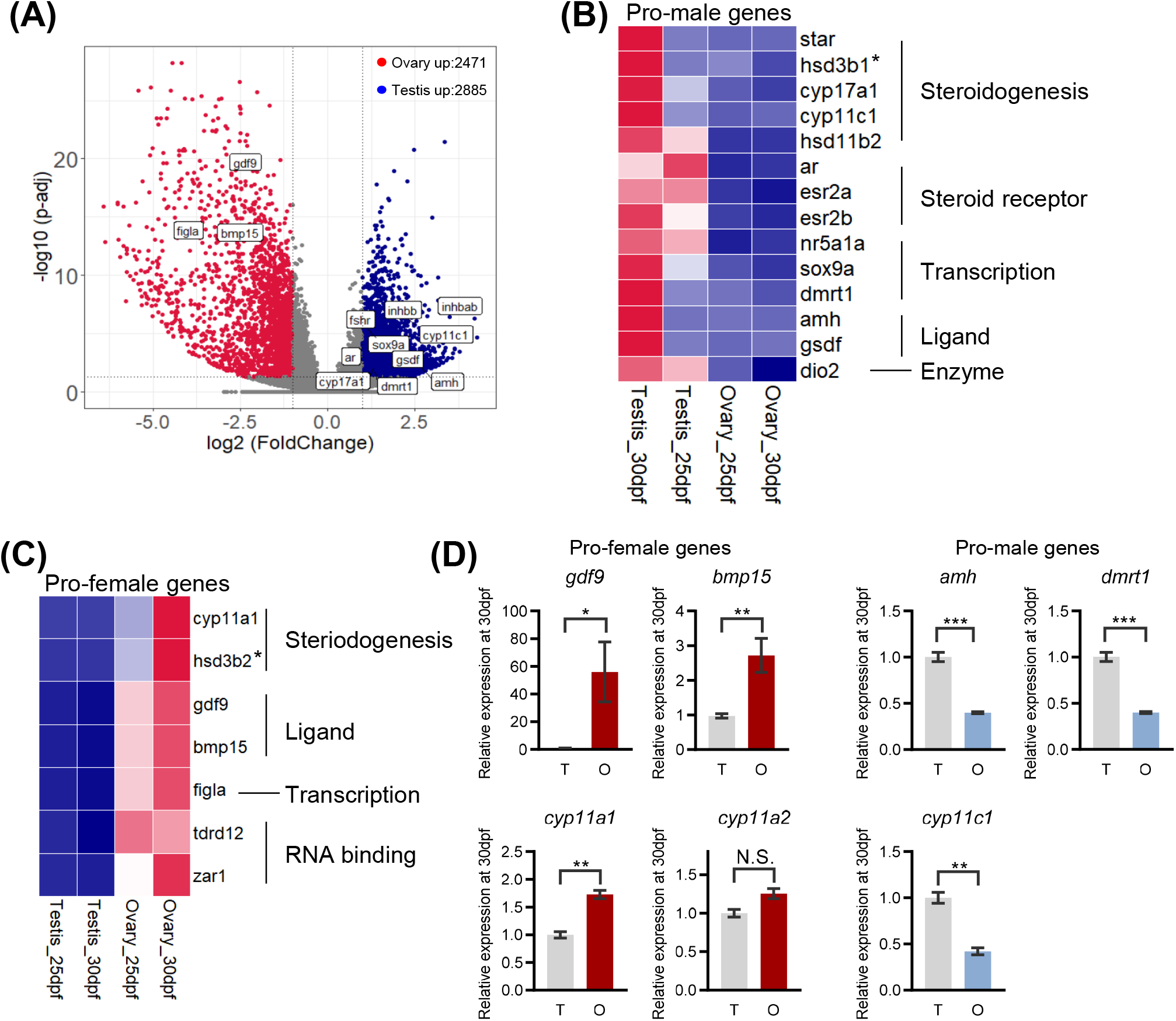
Expression heatmap of selected genes associated with gonad development and RT-qPCR verification. (A) The volcano plot showing DEGs between the presumptive testis and the presumptive ovary. (B) The expression heatmap of the selected DEGs that were known to be involved in gonad development. (C-F) The RT-qPCR analysis of gdf9 (B), bmp15 (C), cyp11a1 (D), amh (E), dmrt1 (F) on the testes and ovaries at 30 dpf, n=3 replicates of biological repeats.

### 3.3 Ovary-enriched biological processes and cellular components

To know what happened to the ovary at the onset of its differentiation, we carried out GO enrichment analysis for biological process and cellular component using the list of the ovary up-regulated genes (Table S3). The representative biological processes and cellular components were shown in Figure 3 A-D. Besides, a GO enrichment map network after removing redundant terms was obtained to globally view the potential functional modules and their relationship (Figure 3 E). Several differentially enriched biological processes and cellular components were listed and discussed below.

**Fig. 3.**
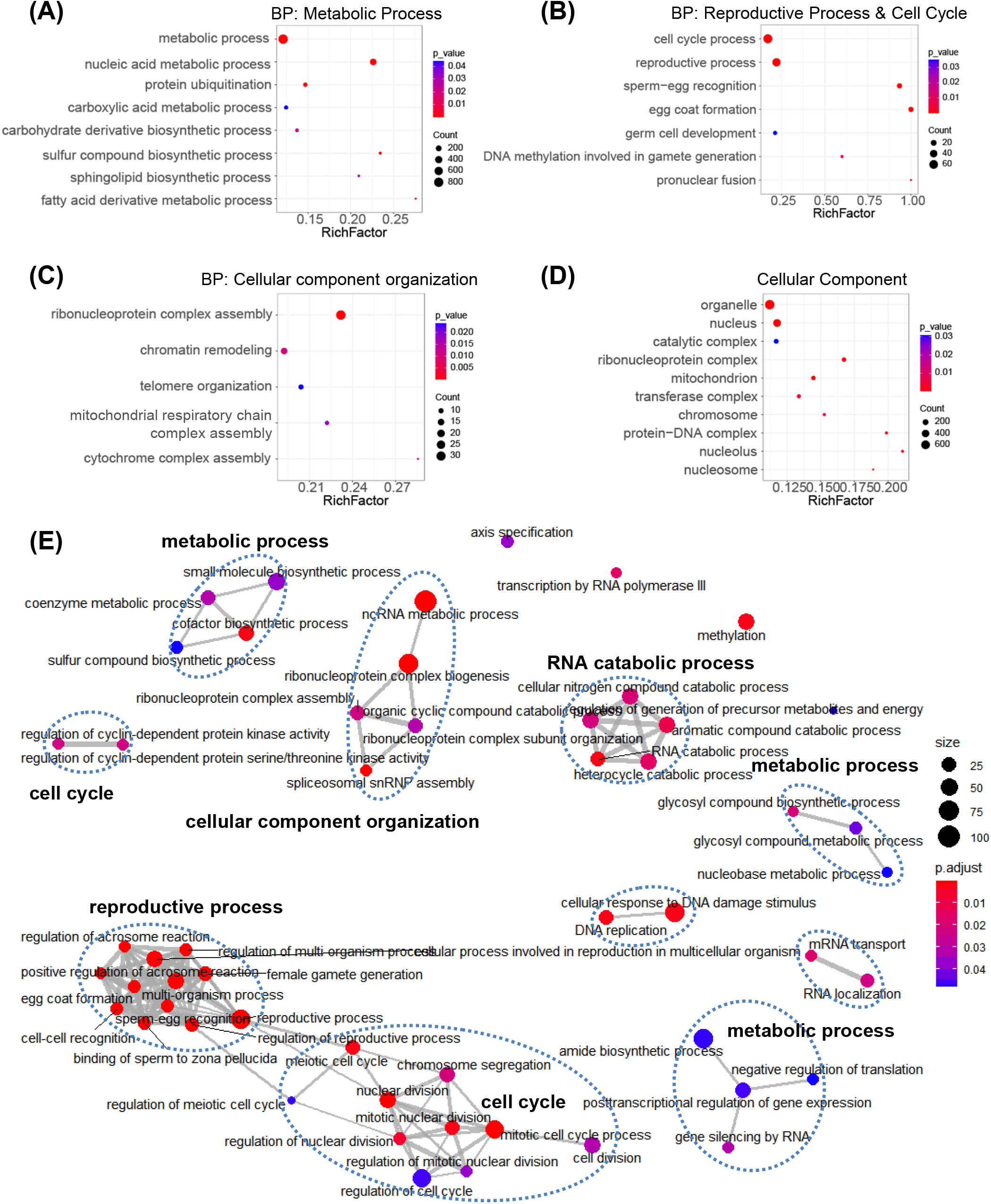
The ovary-enriched biological processes and cellular components at the onset of ovary differentiation. (A) The representative GO terms showing the ovary-enriched metabolic-related biological processes. (B) The representative GO terms showing the ovary-enriched reproductive and cell cycle-related biological processes. (C) The representative GO terms showing the ovary-enriched cellular component organization-related biological processes. (D) The representative GO terms showing the ovary-enriched cellular components. (E) The simplified GO network showing the ovary-enriched biological processes.

#### Maternal synthesis

A class of enriched biological processes are the metabolic processes of different substances including nucleic acid (such as genomic DNA, mitochondrial DNA, mRNA, rRNA, tRNA and non-coding RNA (ncRNA)), protein (the carboxylic acid metabolic process and the sulfur compound biosynthetic process are both related to the biosynthesis of amino acid), carbohydrate derivatives and fatty acid derivatives (Figure 3 A). Those different substances could be the maternal materials which are essential for supporting the embryonic development (Brooks, Tyler et al. 1997). Thus, our data support the view that the active synthesis of various substances is an iconic event in the ovary, and the time when an ovary synthesizes maternal materials is from the onset of differentiation.

#### DNA repair and high transcription activity

The enriched process metabolism of nucleic acid implied that the ovaries were actively generating RNA transcripts at the differentiation period. This is consistent with our observation on the nuclear appearance of oocyte I which has a large cell nucleus with loose chromatin (Ye, Zhu et al. 2019). The enriched process of cellular response to DNA damage stimulus indicated a process of DNA repair which may be related to high transcription activity, as RNA transcription is a major threat to the genome stability and causes DNA double-strand breaks (Figure 3E) (Aguilera and Garcia-Muse 2012, Marnef, Cohen et al. 2017). The enriched process of ncRNA metabolism would be involved in the DNA repair because ncRNA are major regulators of the DNA repair (Sharma and Misteli 2013, Thapar 2018).

#### RNA degradation

It was noted that RNA catabolic process was also enriched, suggesting the machinery for RNA degradation is active (Figure 3 E), and this machinery may be related to the genetic robustness. The genetic robustness is the genetic buffering system that many organisms develop to maintain normal development in the presence of genetic mutants. One of the mechanisms for genetic robustness is the genetic compensation response which is mediated by non-sense mRNA mediated decay/degradation (Ma, Zhu et al. 2019). Thus, we hypothesized that the active RNA degradation machinery along with the increased chromatin accessibility provide an ideal intracellular environment for the genetic robustness, and the genetic compensation response would likely happen in the early-stage oocytes during the ovary differentiation in a gene-mutated zebrafish.

#### Protein ubiquitination and sphingolipid biosynthesis

Other biological processes related to ubiquitination and sphingolipid biosynthesis were also enriched (Figure 3 A). On one hand, protein ubiquitination facilitates protein degradation, and in mammals, this process regulates the cell cycle, fertilization and oocyte maturation (Wu, Li et al. 2021). The genes involved in ubiquitination are thought to be related to the egg quality in Sea Bass (Żarski, Nguyen et al. 2017). Therefore, our data supported a hypothesis that protein ubiquitination might play a conserved role for the normal development of zebrafish oocyte as in the mammals. On the other hand, sphingolipid is an essential structural components of cell membrane. From stage I to maturity, a zebrafish oocyte increases in diameter from 7 microns to 750 microns with the cell surface area increasing over 10,000 times (Selman, Wallace et al. 1993). Thus, it is expected to see the phospholipid biosynthesis is active in our data, which would provide the material for cell membrane enlargement.

#### Reproductive process and cell cycle

The reproductive process and those biological processes relative to sperm-egg recognition such as egg coat formation and pronuclear fusion were enriched (Figure 3 B, E). This data showed a unique character of a female gamete - the fertilization ability, and an oocyte may start to prepare for this competence at the beginning of its differentiation. The cell cycle process including meiosis and mitosis was also enriched (Figure 3 B, E), which might be related to the meiotic arrest of oocyte and the cell proliferation of gonad somatic cells at these stages.

#### Cellular components and their organization

Several cellular components (ribonucleoprotein complex, chromatin, telomere and mitochondrial component) were actively organized, and those components were also displayed in the list of ovary-enriched cellular components (Figure 3 C, D). The organization of cellular components is adaptive with the enriched biological processes. For example, the organization of ribonucleoprotein complex is related to the spliceosomal snRNP assembly for RNA post-transcriptional processing (Figure 3E), and the chromatin remodeling is highly related to the active transcription event discussed above. Mitochondria are responsible for the energy production of cells, and are essential for optimal oocyte maturation, fertilization and early embryonic development (Babayev and Seli 2015). Mitochondria of a new embryos is not from the sperm but from the oocyte (May-Panloup, Chretien et al. 2007). The mitochondrial DNA copy number of an oocyte is about 100 times as much as that of a somatic cell in bovine (Michaels, Hauswirth et al. 1982), and it is considered as a marker of oocyte quality in mice (Chiaratti and Meirelles 2010). In our data, two mitochondrial associated processes were enriched, suggesting that the ovary development may be a high-energy consuming process. As an in-vitro developed species, the zebrafish embryonic development might highly rely on the copy number of mitochondria from a fish egg.

In addition, the ovary up-regulated genes were not enriched in any growth factor or morphogenic protein-mediated signaling pathway. One possible reason would be that the signaling pathway was contributed by minor cells whose transcriptional contribution would be diluted by the bulk-seq data of the whole ovary. This limitation could be addressed by single-cell sequencing in the future. Taken together, the ovarian GO analysis drew a picture of differentiating ovary in which processes about maternal synthesis, DNA repair, RNA transcription, RNA degradation, protein ubiquitination, sphingolipid biosynthesis, reproduction, cell proliferation, cellular components organization are active.

### 3.4 Testis-enriched biological processes, cellular components and signaling pathways

To know what happened to the testis at the onset of its differentiation, we also performed GO analysis using the testis up-regulated genes (Table S4). Ten major biological processes represented the characteristics of the differentiating testis (Figure 4A). In contrast to the metabolic character of ovary, the major character of testis is the developmental characteristic including morphogenesis, growth, cell differentiation and proliferation (Figure 4A). The locomotion and biological adhesion are both highly relative to morphogenesis (Figure 4A). The enriched processes about rhythm showed another characteristics of the differentiating testis (Figure 4A). The rhythmic functions in testis is thought to rely on the periodicity of seminiferous epithelium and Leydig cell (Bittman 2016). The former regulates the timing of spermatogenesis through retinoic acid, and the later regulates the output of the steroid hormones (Bittman 2016). Our data showed the rhythm process enriched and the rhythm-related genes up-regulated in the zebrafish testes at the onset of their differentiation, supporting the view that testis is a rhythm organ. Moreover, the immune system process was enriched, implying a critical role of immunoregulation during the testis differentiation which will be further analyzed below. The testis-enriched cellular components revealed the active elements on extracellular region, cell membrane, and chromatin (Figure 4B), which accommodates the developmental changes of differentiating testis.

**Fig 4.**
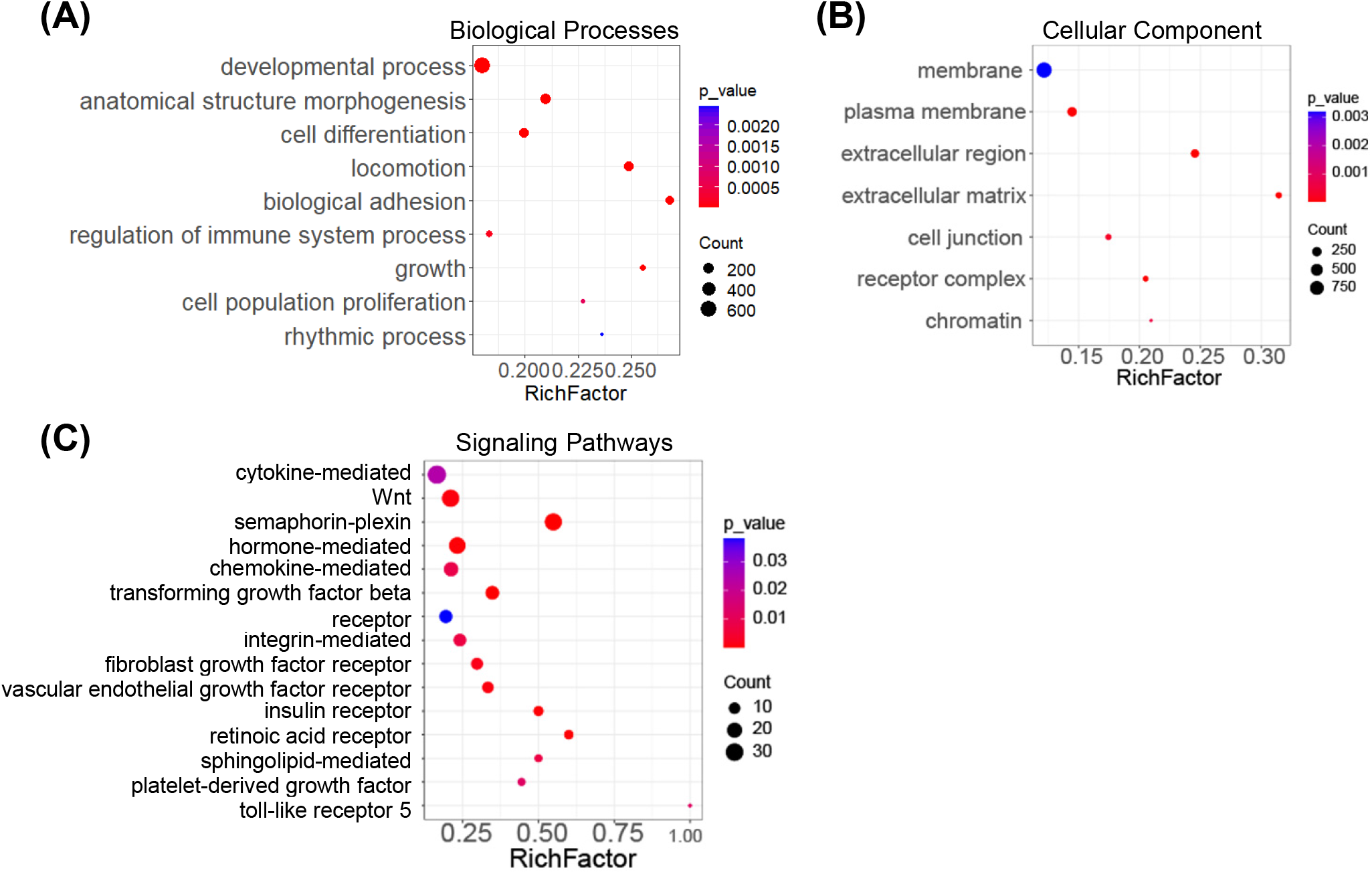
The testis-enriched biological processes, cellular components and signaling pathways at the onset of testis differentiation. (A) The representative GO terms on the testis-enriched biological processes. (B) The representative GO terms on the testis-enriched cellular components. (C) The representative GO terms on the testis-enriched signaling pathways.

Because the testis differentiation showed developmental properties, we then investigated what growth factor - mediated signaling pathways were involved. We found that the testis up-regulated genes were enriched into fifteen signaling pathways (Figure 4C), and the genes related to each pathway were listed as Figure 5. Among those, signaling pathways relative to Wnt (Dong, Tan et al. 2015), hormone (Sofikitis, Giotitsas et al. 2008), transforming growth factor beta (Young, Wakitani et al. 2015), Notch (Tang, Brennan et al. 2008, Ng, Qian et al. 2019), integrin (Kanatsu-Shinohara, Takehashi et al. 2008), fibroblast growth factor (Jiang, Skibba et al. 2013), insulin (Reinecke 2010), vascular endothelial growth factor (Sargent, McFee et al. 2015), retinoic acid (Endo, Mikedis et al. 2019), sphingolipid (Tilly and Kolesnick 1999), platelet-derived growth factor (Basciani, Mariani et al. 2010), Toll-like receptor (Shang, Zhang et al. 2011) were known to play critical roles in the testis development, and those signaling pathways might play conserved roles during the testis differentiation in zebrafish. However, to our knowledge, there is no report about the function of semaphorin-plexin signaling pathway in the testis development, and its function need to be experimentally investigated in the future. The cytokine-mediated and chemokine-mediated signaling pathway may be related the immunoregulation which will be discussed below.

**Fig. 5.**
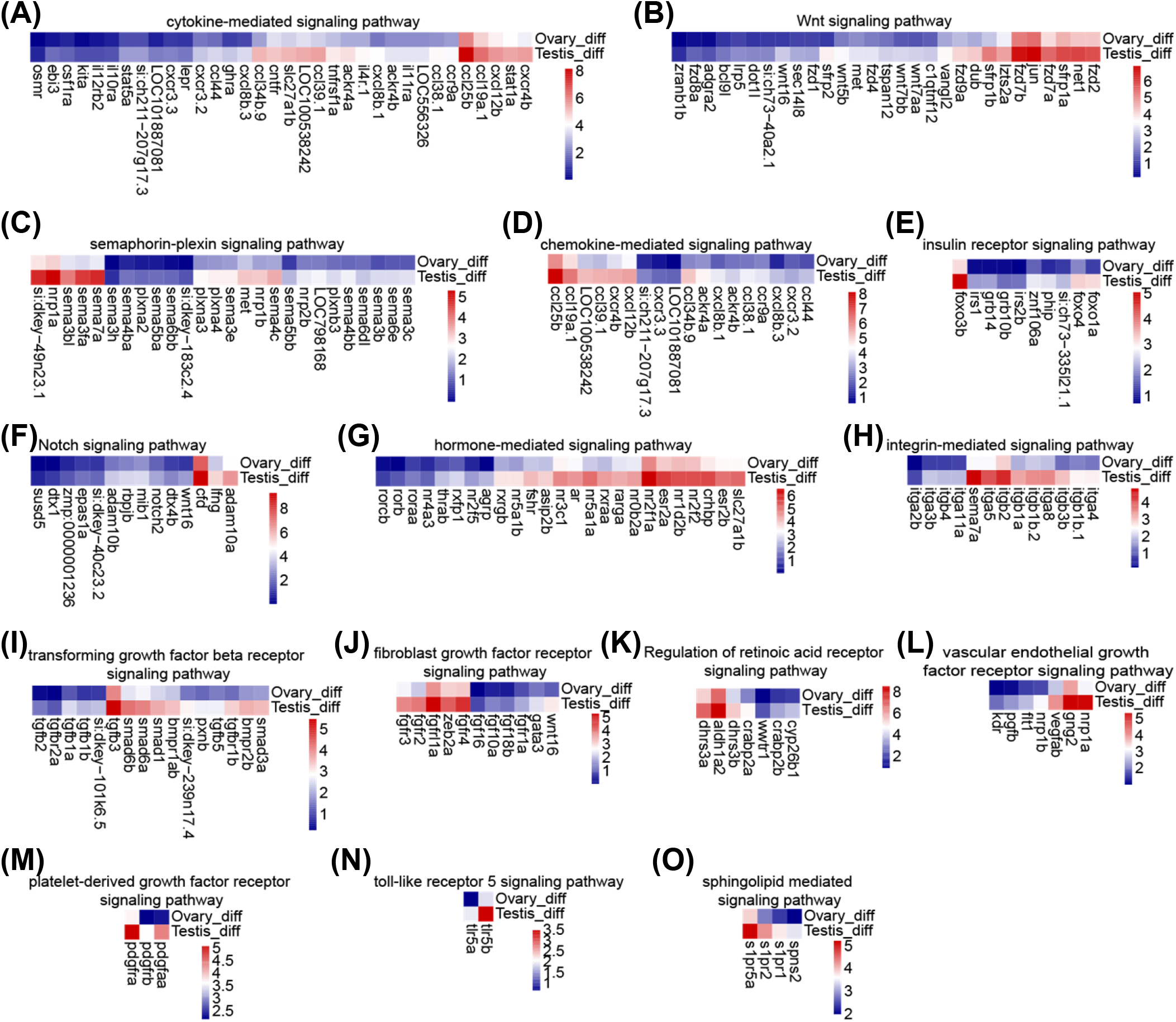
The expression heatmap of the genes for each signaling pathways in the Fig. 4C. (A) Cytokine-mediated signaling pathway; (B) Wnt signaling pathway; (C) Semaphorin-plexin signaling; (D) Chemokine-mediated signaling pathway; (E) Insulin receptor signaling pathway; (F) Notch signaling pathway; (G) Hormone-mediated signaling pathway; (H) Integrin-mediated signaling pathway; (I) Transforming growth factor beta receptor signaling pathway; (J) Fibroblast growth factor receptor signaling pathway; (K) Regulation of retinoic acid receptor signaling pathway; (L) Vascular endothelial growth factor receptor signaling pathway; (M) Platelet-derived growth factor receptor signaling pathway; (N) Toll-like receptor 5 signaling pathway; (O) Sphingolipid mediated signaling pathway.

### 3.5 The immunoregulation in the testis differentiation

In recent studies, we found that the oocytes were undergoing apoptosis in the juvenile testis at 30 dpf, resulting in immune response and inflammation (Ye, Zhu et al. 2019, Li, Zhang et al. 2020). Excessive inflammation in the testis would disrupt both germ cells and gonadal somatic cells leading to male infertility, and the regulatory T cells suppress immune response to a reasonable level and maintain the immune homeostasis in testis (Li, Zhang et al. 2020). The number of immune-related genes in the testis up-regulated DEGs was much more than that in the ovary (Figure 6A), indicating a higher activity of immune response in the testis than that in the ovary at the onset of gonad differentiation.

**Fig. 6.**
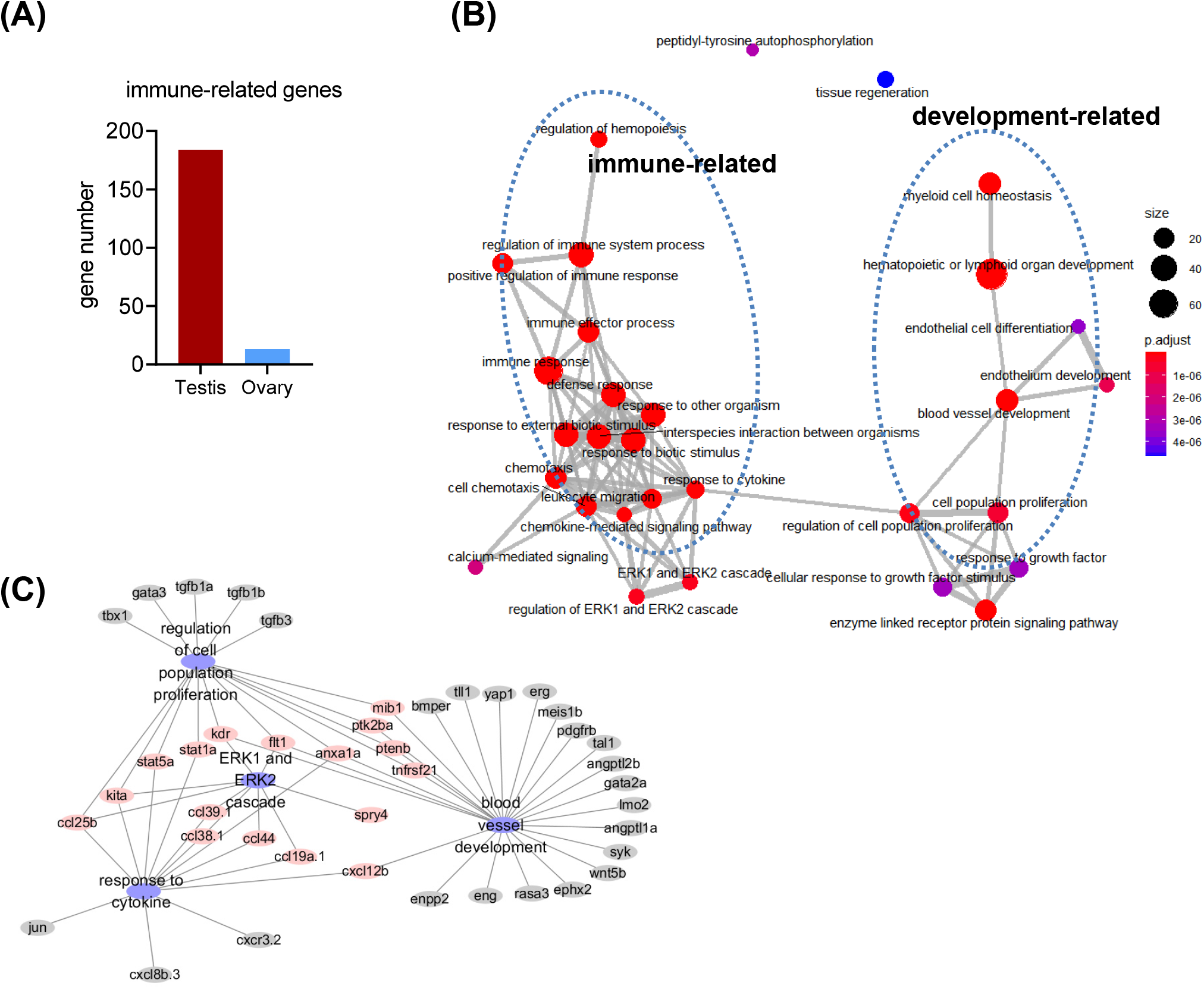
The immunoregulation in the testis differentiation. (A) The number of immune-related gene in the testis up-regulated genes and the ovary up-regulated genes. (B) The simplified GO network showing the relationship between immune-related processes and development-related process. (C) The category gene network showing the genes involved in the cytokine-induced cell proliferation and vasculogenesis.

The testis immune-related genes were then used for the GO enrichment analysis and the simplified GO network was visualized (Figure 6B). The GO network showed two clusters of interrelated terms. One was the immune-related processes and the other was development-related processes (Figure 6B). In the cluster of immune-related processes, the ERK1 and ERK2 cascade was found to be high relative to the cytokine-mediated signaling pathway (Figure 6B), and the two clusters were connected through two nodes-response to cytokine and regulation of cell population proliferation (Figure 6B), implying the immune-related processes might promote the testis differentiation via regulating the cell proliferation.

Moreover, in the cluster of development-related processes, regulation of cell population proliferation was linked to blood vessel development (Figure 6B). In mammals and human, it was proposed that the cytokines and the ERK1/2 signaling pathway are likely involved in Sertoli cell proliferation (Meroni, Galardo et al. 2019), and the progression of chronic testicular inflammation is associated with the proliferation of testicular endothelial cells in rodents (Gualdoni, Jacobo et al. 2021). Although little is known about the blood vessel formation in the testis under normal physiological and genetic condition, our data support a hypothesis that the vasculogenesis might begin at the onset of testis differentiation and might be promoted by immune-induced cytokines production. We further plotted the category gene network to show the specific genes involved in the immune-induced cell proliferation and vasculogenesis in the testis (Figure 6C). The highlighted genes linking different processed need to be experimentally proved in the future. Taken together, these data imply a novel role of immunoregulation in vasculogenesis during testis development. Functional studies of the newly proposed roles of immunoregulation in testis differentiation are needed and will provide valuable insights into gonad development in fish.

## 4. Conclusions

In this study, we have provided the first transcriptional landscape of ovary and testis at the onset of differentiation by utilizing intact juvenile gonads, which greatly expand our knowledge of the gonad differentiation in zebrafish. The sexually dimorphic expression appears at around 25 dpf in zebrafish. During ovary differentiation, the biological processes related to maternal synthesis, DNA repair, RNA transcription, RNA degradation, protein ubiquitination, sphingolipid biosynthesis, reproductive, cell cycle, and cellular components organization were enriched. During testis differentiation, the biological processes related to morphogenesis, growth, cell differentiation, cell proliferation, locomotion, biological adhesion, rhythm, and immunoregulation were highly enriched. Fifteen signaling pathways and related genes were found to be involved in the testis differentiation. Notably, our study suggests that the immune-related processes are extensively involved in the regulation of testis development. Overall, this study provides novel insights into gonad differentiation in zebrafish.

## 5. Ethical Statement

All animal procedures performed in this research were in accordance with the ethical standards of the Institutional Animal Care and Use Committee of the Institute of Hydrobiology, Chinese Academy of Sciences.

## 6. Declaration of competing Interest

The authors declare that they have no known competing financial interests or personal relationships that could have appeared to influence the work reported in this paper.

## 6. Funding

This study was supported by grants from National Natural Science Foundation of China (31872550, 31721005, and 31871305), National Key R&D Program of China (2018YFD0901205), Strategic Priority Research Program of the Chinese Academy of Sciences (XDA24010108), State Key Laboratory of Freshwater Ecology and Biotechnology (2019FBZ05, 2020FB08), Fundamental Research Funds for the Central Universities (2662019PY003, 2662020PY001), HZAU-AGIS Cooperation Fund (SZYJY2021010), and Huazhong Agricultural University Scientific & Technological Self-innovation Foundation (2016RC011). The funding body had no roles in the design of the study and collection, analysis, and interpretation of data and in writing the manuscript.

## 8. Acknowledgments

We thank Kuoyu Li from China Zebrafish Resource Center for many helpful discussions about zebrafish breeding and fish health. We thank Zhixian Qiao and Xiaocui Chai from Analysis and Testing Center of Institute of Hydrobiology for technique support of high-throughput sequencing.

## 9. Author Contributions

Ding Ye: concept and design of the original research, acquisition of data, interpretation of data, draft of the article; Yi-Xuan Tu: analysis and visualization of data; Houpeng Wang: acquisition of data; Mudan He: acquisition of data; Yaqing Wang: acquisition of data; Zhengfang Chen: acquisition of data; Yonghua Sun: critical revision of intellectual content, revision and finalization of the manuscript; Zhen-Xia Chen: critical revision of intellectual content.

## 10. Supplementary Materials

Table S1: The primer list for the RT-qPCR analysis

Table S2: Differentially expressed gene list

Table S3: GO enrichment result of ovary up-regulated DEGs

Table S4: GO enrichment result of testis up-regulated DEGs

## Notes

### Competing Interest Statement

The authors have declared no competing interest.

